# A PATHOGENIC NFKB2 VARIANT CAUSES COMBINED IMMUNODEFICIENCY WITH THYMIC AND PERIPHERAL LYMPHOID DEFECTS IN RATS

**DOI:** 10.64898/2026.10.02.756202

**Authors:** Séverine Remy, Laurent Tesson, Claire Usal, Marie Massoda, Séverine Ménoret, Irina Proekt, David Yu, Axel You, Matthieu Giraud, Mark S. Anderson, Ignacio Anegon

## Abstract

The non-canonical NF-κB pathway, in which NFKB2 plays a central role, is critical for lymphoid organ development and immune homeostasis. Pathogenic NFKB2 variants have been strongly associated with a broad spectrum of human pathologies, notably immunodysregulation syndromes, characterized by recurrent infections, autoantibody production and endocrine abnormalities. The mechanisms underlying the resulting immune defects remain incompletely understood. In the present study, we characterized the immune phenotype of a novel rat model harboring a *Nfkb2* non-sense mutation located in the c-terminal region of the gene (c.2602T>A; p.Tyr867*, *Nfkb2^p.Tyr867*^*). This variant has been identified in patients with a familial form of common variable immunodeficiency (CVID) presenting with autoimmunity manifestations. Using this model, we demonstrate that mutated NFKB2 resulted in profound alterations of the thymic microenvironment, including reduced K5^+^ medullary thymic epithelial cells and decreased AIRE expression, accompanied by altered thymocyte development and impaired thymic regulatory T populations. Peripheral lymphoid organization was also profoundly affected, with absence of lymph nodes, defects in the splenic marginal zone, and alterations of CD4^+^ and CD8^+^ T-cell homeostasis, including impaired Treg populations and reduced ICOS expression on CD4^+^ T cells. The B-cell compartment was markedly altered, with impaired B-cell maturation, characterized by accumulation of IgM^+^IgD^−^ transitional cells and a marked reduction of IgD^+^ mature B cells. These abnormalities were associated with impaired humoral immunity and complete absence of circulating immunoglobulins.

Collectively, our findings define a systemic combined immunodeficiency phenotype in the Nfkb2-mutated rat, highlighting its utility for studying the role of non-canonical NF-kB signaling in immune system organization and adaptative immunity.

## Introduction

The non-canonical NF-κB pathway relies on the p100/p52 protein encoded by *NFKB2*, which plays a central role in immune homeostasis. In contrast to canonical NF-κB signaling, that it is mainly activated by TNFα, this pathway is selectively activated downstream of receptors, including CD40, Receptor Activator of Nuclear Factor-κB (RANK), Lymphotoxin beta receptor (LT-βR) and BAFF receptor (BAFF-R), and plays a critical role in B-cell maturation, lymphoid organogenesis, thymic epithelial cell differentiation and immune tolerance. NF-κB2 signaling has also been implicated in the development and functional regulation of the anterior pituitary, suggesting a molecular link between immune and endocrine homeostasis. Germline heterozygous mutations in *NFKB2* are associated with a spectrum of human immunopathologies, most notably common variable immunodeficiency (CVID) and DAVID syndrome (Deficient Anterior pituitary with Variable Immune Deficiency). These disorders are characterized by hypogammaglobulinemia, recurrent infections, autoimmunity and variable endocrine dysfunctions affecting anterior pituitary hormones [1–5].

Pathogenic variants cluster in the C-terminal region of *NFKB2* and disrupt NF-κB-inducing kinase (NIK) dependent processing of the p100 precursor into the transcriptionally active p52 subunit. Mechanistically, C-terminal *NFKB2* variants impair p100 phosphorylation and proteasomal processing, resulting in the accumulation of an unprocessable inhibitory p100. This aberrant protein acts as a dominant negative regulator, attenuating non-canonical NF-κB signaling and partially interfering with canonical NF-κB activation. The resulting p52 insufficiency compromises B-cell survival, thymic T-cell maturation, regulatory T-cell homeostasis and central tolerance.

Study of mouse models targeting components of the non-canonical NF-kB pathway, including *Nfkb2^−/-^* (lacking p100/p52) [6,7]*, p52-deficient mice* [8]*, Rel-B*-deficient mice [9], *Nfkb2^Lym/Lym1^* mice (same mutation in p100as in this rat model) [10], and *aly/aly* mice (point mutation in NIK) [11,12], have revealed profound defects in lymphoid organ organization and immune function. Several of these models exhibit variable degrees of thymic medullary disruption [9,10,12–16], impaired mTEC development [14,17], reduced thymic expression of the autoimmune regulator AIRE [7,10], together with lymphocyte dysregulation and inflammatory [7,9–11,16] or autoimmune manifestations, including multiorgan lymphocytic infiltration [7,9,12–16] and, in some models, autoantibodies production [7,16–18].

Importantly, studies of distinct NFKB2 variants have demonstrated that different molecular alterations can result in markedly different immune phenotypes, depending on their effects on p100 processing and p52 production, highlighting the importance of the specific consequences of individual pathogenic variants NFKB2 [16].

Rats represent a relevant translational model for studying immune-endocrine interactions, as their immune architecture and pituitary physiology are closer to those of humans than those of mice. Indeed, several rat models display immune and endocrine phenotypes that more recapitulate human disease [19]. Our group has previously generated and characterized an AIRE-deficient rat model that develops spontaneous multi-organ autoimmunity and circulating auto-antibodies, closely mirroring the clinical spectrum of Autoimmune Polyendocrinopathy- Candidiasis-Ectodermal Dystrophy (APECED) patients [20]. In contrast to AIRE-deficient mice, these rats exhibit a broader and more penetrant autoimmune phenotype [20], underscoring the value of the rat model for investigating mechanisms of central tolerance.

The observations reported in *Nfkb2*-mutant mice, including defective mTEC development and reduced AIRE expression [7,17], suggest that NF-κB2 signaling may intersect with AIRE- dependent transcriptional programs in thymic epithelial cells.

To date, no *Nfkb2*-deficient or mutated rat model has been reported. Here, we report the generation of a rat model carrying the pathogenic *p.Tyr867* Nfkb2* mutation. The premature stop codon caused by this mutation generates a NFKB2 p100 precursor protein truncated at the C-terminal phosphorylation site, which disrupts the p100 precursor processing into the transcriptionally active p52 subunit. This results in the accumulation of an unprocessable inhibitory p100 that acts as a dominant negative regulator, attenuating non-canonical and canonical NF-κB signaling. This model provides an in vivo system to investigate NFKB2- dependent signaling in immune homeostasis and central tolerance. We show that *Nfkb2^p.Tyr867*^* mutation results in a profound combined immunological defect affecting lymphoid organ architecture and adaptative immune responses, in the absence of overt autoimmunity or endocrine abnormalities.

## Results

### P100 processing defect in *Nfkb2^p.Tyr867*^* mutant rats

To investigate the involvement of the non-canonical NF-κB pathway in the regulation of immune and inflammatory responses, we generated a novel rat line carrying the c.2598C>A (p.Tyr867*) mutation, mimicking the mutation described in the *Nfkb2^Lym1/Lym1^* mice [10] and the pathogenic variant in patients with DAVID syndrome [1]. The mutation was introduced by electroporation into the rat zygote [21] using a single-guide RNA designed to target exon 21 of the rat *Nfkb2* gene and a 100nt ssODN including a stop codon at position Y867, an EcoRI restriction site, and other point mutations to avoid CRISPR cleavage after donor DNA insertion **(Fig.1A)**. Two rat founders carrying the expected mutation were chosen to derive two distinct lines to confirm the phenotype. In a representative cohort, the p.Tyr867* mutation was transmitted in one rat (named “c”) out of seven offspring and three rats (named “i, j, m”) out of six offspring, issued from mating of the 1.3 and 2.4 founder rats, respectively, with wild-type (WT) rats **(Fig.1B)**. *EcoRI* digestion suggested a heterozygous Knock-In (KI) insertion in these four animals, given the size of r*Nfkb2* PCR products (528pb, WT allele; 335bp and 193bp, KI allele) **(Fig.1B)**. The genotype of these four rats was confirmed by sequencing PCR products (**Fig.1C**).

**Figure 1.**
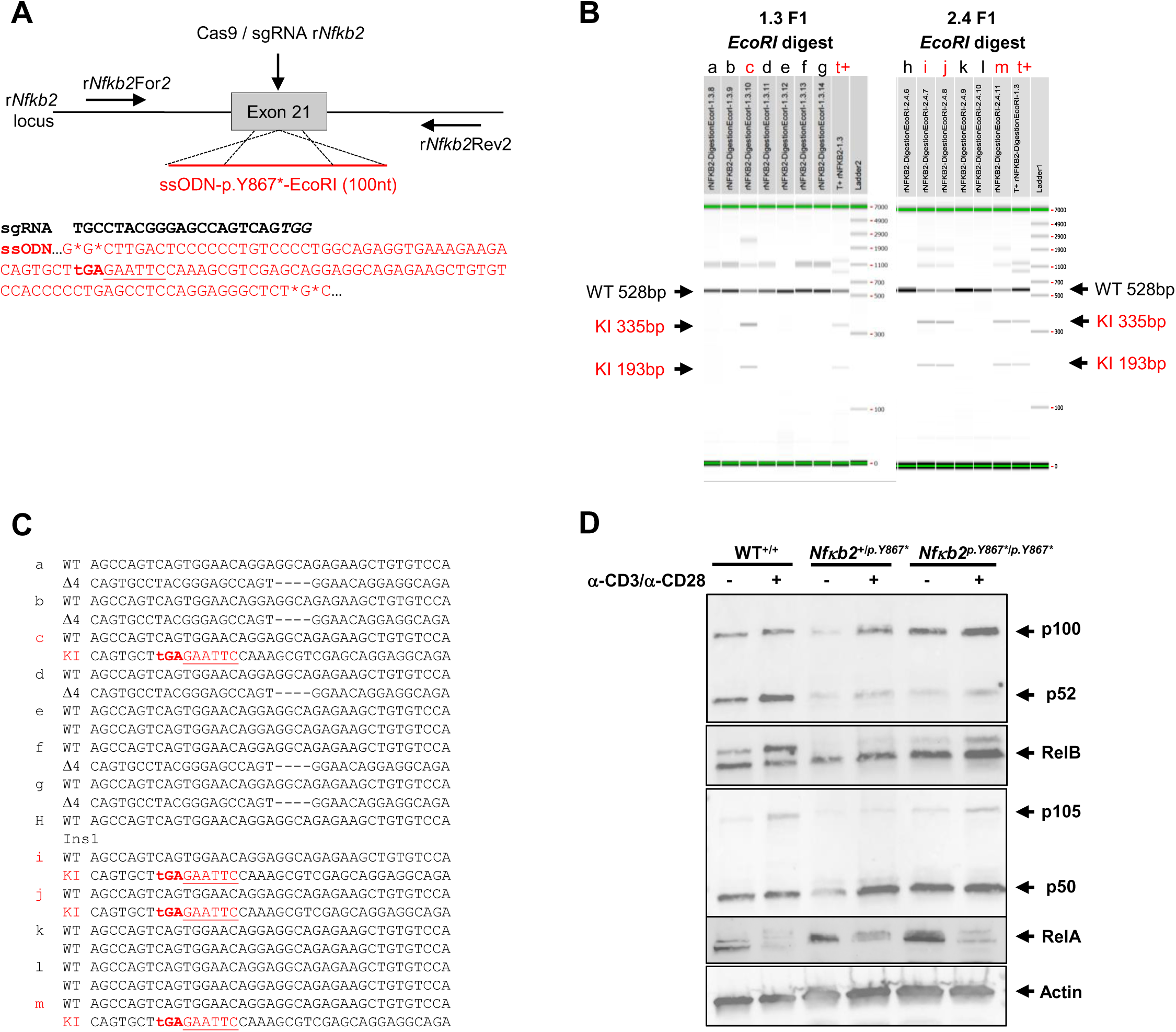
Generation of p.Tyr867*-NFkB2 mutant rats. **(A)** Schematic illustration of the target sequence (+PAM in italic) in exon 21 of the rat *Nfkb2* gene and the 100 nt ssODN designed to insert a stop codon (tGA, bold sequence) and an *EcoRI* restriction site (GAATTC, red underlined sequence to facilitate genotyping. Additional mutations were introduced into ssODN to prevent cleavage of this template (indicated by stars). The position of each primer (r*Nfkb2*For2 and r*Nfkb2*Rev2) is indicated on the schematic rat *Nfkb2* locus. **(B)** *Nfkb2* PCR analysis of a representative panel of 7 (a, b, c, d, e, f, g) and 6 (h, i, j, k, l, m) rats from the F1 generation of lines 1.3 and 2.4 respectively. Capillary gels show the r*Nfkb2* PCR products of the 528 bp expected size, and 335 bp and 193 bp after *EcoRI* digestion. One animal (c) of the 1.3 line and three (i, j, m) of the 2.4 line showed *EcoRI* digestion suggesting heterozygous KI insertion. **(C)** Sequencing of the subcloned PCR products from positive animals F1 HE animals. Deletions are indicated by “D”, insertions by “Ins” and knock-in events are indicated by “KI”. *EcoRI* recognition sites are underlined in inserted alleles. In six animals (a, b, d, f, g, h), NHEJ events were observed. Four animals are heterozygous KI (c, i, j, m) and three are WT (e, k, l). All animals have one chromosome WT. **(D)** Representative western blot analysis of p100/p52, RelB, p105/p50, RelA expression in lysates of WT^+/+^, *Nfkb2^+/p.Y867*^* or *Nfkb2^p.Y867*/p.Y867*^*splenocytes overnight stimulated with plate-bound anti-CD3 plus anti-CD28. Actin was used as loading control.

The premature stop codon caused by the mutation p.Tyr867* is predicted to result in an NFKB2 protein truncated at the C-terminal phosphorylation site with a predicted mass of 93.618 kDa, compared to the native form (MW: 97.134 kDa). Although western blot analysis cannot determine this difference in size, we used it to analyze the impact of the p.Y867*mutation on the expression of proteins involved in the NF-κB2 non-canonical pathway (p100/p52, RelB) and in the NF-κB1 canonical pathway (p105/p50, RelA, c-Rel). An increase of p100 was observed in WT, heterozygous (HE) or homozygous (HO) splenocytes activated with anti-CD23 and anti-CD28 antibodies, while p52 subunit was weakly detected in HE and HO splenocytes, suggesting a p100 processing defect **(Fig. 1D)**. The accumulation of p100 observed in HO lysates seems to corroborate this hypothesis. In contrast, the total protein levels of RelB, p105/p50 and RelA remained unchanged by the p.Y867* mutation, suggesting that the defect primarily affects p100 processing rather than the overall abundance of other NF-kB family members **(Fig. 1D)**.

Together, these results confirm the generation of 2 lines of ***Nfkb2^p.Tyr867*^***rats with predominant impaired activation of the non-canonical NF-κB pathway, without altering the expression of other major NF-κB pathway components.

### Clinical manifestations in *Nfkb2^p.Y867*/ p.Y867*^*rats

Rats carrying the HO p.Y867* mutation showed a reduced lifespan, with a median survival of 217 days (line 2.4) and 367 days (line 1.3) compared to HE rats (median survival: 378 d, line 2.4; 460d, line 1.3) and wild-type (littermate) rats (median survival: 530d) **(Fig. 2A)**.

**Figure 2.**
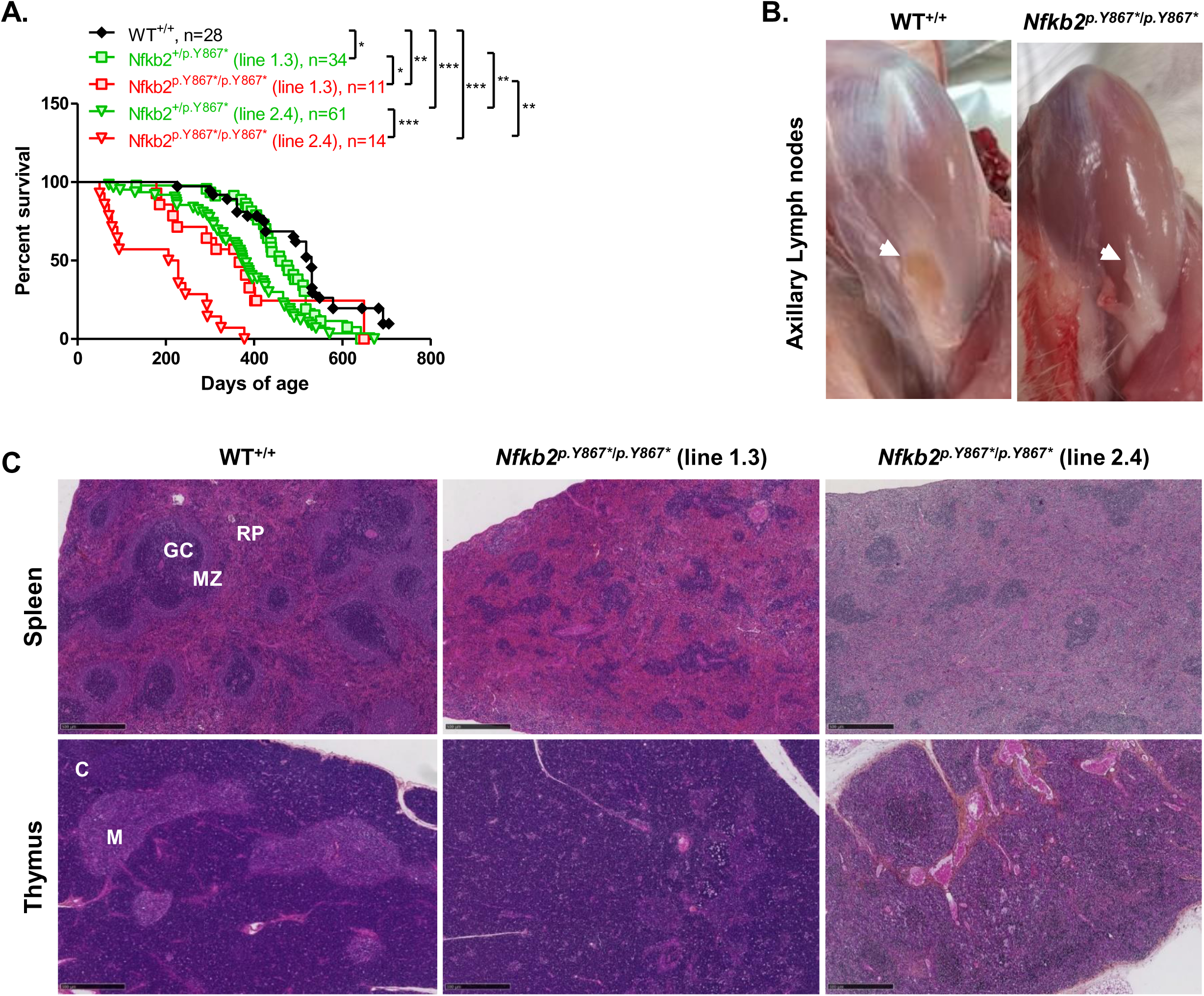
Reduced lifespan and defective lymphoid organogenesis in *Nfkb2^p.Y867*/p.Y867*^*rats. **(A)** Comparison of the lifespan of *Nfkb2^+/p.Y867*^*and *Nfkb2^p.Y867*/p.Y867*^* rats from each mutant line (1.3 and 2.4 lines) with WT^+/+^ littermate rats. Each genotype was compared to the WT group, and the heterozygous and homozygous groups were compared to each other, using a log-rank test; ns, not significant; *, p<0.05; **, p<0.01; ***, p<0.001. **(B)** Absence of axillary lymph node in a representative *Nfkb2^p.Y867*/p.Y867*^* rat (line 1.3) (right image) compared to a WT^+/+^ littermate rat (left image). Axillary lymph node areas are indicated by arrows. **(C)** Representative histology of spleen and thymus section from 3-month-old WT^+/+^ littermate (left panel) and *Nfkb2^p.Y867*/p.Y867*^* rats from each mutant line (line 1.3, middle panel and line 2.4, right panel). Hematoxylin-phloxine-saffron staining. RP, red pulp; GC, germinal center; MZ, marginal zone; C, cortex; M, medulla. Scale bars, 500 µM.

Weekly weight monitoring from weaning to 5 months of age revealed no statistical differences in growth of HE and HO *Nfkb2^p.Y867*^* rats (of both lines, of both sex) compared to wild-type counterparts **(supplemental Figure 1)**.

Post-mortem anatomical analysis revealed the complete absence of peripheral lymph nodes (axillary, popliteal, mesenteric) in HO rats **(Fig. 2B and data not shown)**, while rare mesenteric lymph nodes were observed in HE rats (data not shown). Histopathological analyses revealed infiltration of sinusoids by large amphophilic lymphoblastic cells in mesenteric lymph nodes of young heterozygous rats, compared with the normal appearance of lymph nodes in a wild-type rat (data not shown).

Analysis of the spleen in homozygous rats (both line) revealed a marked disruption of splenic architecture, characterized by the absence of B-cell follicles and the lack of marginal zone **(Fig. 2C, upper panels)**, indicating a profound defect in secondary lymphoid organization. In parallel, examination of the thymus showed a marked disorganization of the thymic medulla in these animals (of both lines) **(Fig. 2C, lower panels)**.

Perivascular infiltration by lymphoblastic cells was observed in liver of young HO **(supplemental Figure 2)** and HE rats (data not shown) (of both lines), while no inflammatory infiltrates were detected in in any of the other tissues analyzed (colon, kidney, lung) of these rats **(supplemental Figure 2 and data not shown)**. Histopathological analyses of endocrine tissues revealed no anatomical defects in pancreas, adrenal glands, thyroid, salivary glands pituitary and testis of young HO **(supplemental Figure 3)** and HE rats (data not shown) (of both lines). A reduced, but not significant, pancreatic mass was observed in one-year-old HE and HO rats compared to WT rats, suggesting exocrine pancreatic insufficiency (data not shown).We observed sporadic cases of absence of mature follicles in ovaries of young HO rats (both lines), mammary (line 1.3) or ovarian (both lines) tumors in HE and HO, and dental malocclusion (both lines) or limb bone malformation (line 2.4) (data not shown).

Analysis of biochemical parameters (sodium, potassium, phosphorus, total calcium, creatinine, total protein, glucose, lipase and alkaline phosphatases) in the serum of 3- and 7- month-old HO rats (both lines) revealed no significant differences compared to control rats **(supplemental Figure 4)**. A significant decrease in plasma albumin was observed in both young and old rats of line 2.4, while a significant increase in transaminases was detected in older animals of lines 2.4 (ASAT/ALAT) and 1.3 (ALAT), suggesting liver damage **(supplemental Figure 4)**. These data combined with the inflammatory foci observed in the in liver at 3 months, suggest progressive inflammatory hepatocellular damage.

The analysis of blood platelets, red blood cells, hemoglobin, hematocrit, total leucocytes including lymphocytes, monocytes, neutrophils, eosinophils and granulocytes did not show differences with controls (data not shown).

Adrenocorticotropic hormone (ACTH), thyroid-stimulating hormone (TSH), growth hormone (GH), follicle-stimulating hormone (FSH), luteinizing hormone (LH) and prolactin levels in homozygous rats of lines 1.3 and 2.4 did not differ significantly from those of wild-type rats, with the exception of a 1.5-fold increase in ACTH levels in young homozygotes rats of line 2.4 **(supplemental Figure 5)**, suggesting activation or dysregulation of the hypothalamic– pituitary–adrenal (HPA) axis without pituitary histopathological correlation.

Finally, no detectable autoantibodies were observed in *Nfkb2^p.Y867*^*rats. Indeed, no autoantibodies directed against IFNα2 were detected by radioligand binding assay (RLBA) in sera of aged (7 to 14 months) heterozygous (n=53), homozygous (n=11) (both lines combined) and wild-type rats (n=38) **(supplemental Figure 6)**, as they have been detected in humans [22].

Similarly, immunohistochemical analysis of organ-specific autoantibodies, performed on sera of homozygous rats older than 7 months, revealed no reactivity in the colon, ileum, kidney, liver, pancreas or salivary glands (data not shown).

Consequently, the reduced lifespan observed in rats carrying the homozygous *Nfkb2^p.Y867*/ p.Y867*^* mutation is likely multifactorial, resulting from the combined effects of disrupted lymphoid organogenesis and inflammatory damage.

Given the central role of NF-κB2 in lymphoid organogenesis and immune cell development, we next investigated the immunological consequences of the homozygous *Nfkb2^p.Y867*/p.Y867*^* mutation.

### Impaired thymic medulla development in *Nfkb2^p.Y867*/ p.Y867*^*rats

HO mutants exhibited a slight, non-significant reduction in thymic size and weight compared with control and HE rats **(Supplemental figure 7A)**, consistent with a trend toward thymic hypoplasia. Although CD45^+^ cell numbers were preserved **(Supplemental figure 7C)** total thymic cellularity was reduced by 1.5-fold **(Supplemental figure 7B)**, suggesting alterations in the thymic stromal compartment. We therefore examined thymic epithelial cell populations together with thymocyte differentiation.

Homozygous mutants of both *Nfkb2^p.Y867*/ p.Y867*^* lines showed marked disorganization of the thymic medulla **(Fig. 2C)**, together with reduced cytokeratin 5 (CK5) staining and decreased AIRE expression **(Fig. 3A-3B)**, indicating alterations of the medullar thymic epithelial cells (mTEC) compartment and its maturation. Immunologically, these changes were associated with a slight but significant reduction in CD45R^−^ CD4^+^ CD8^+^ double-positive (DP) thymocytes and an increase in the proportion of CD4 single-positive (SP) cells, while CD8 SP thymocytes were unaffected **(Fig.4A-4B)**. These findings are consistent with impaired thymic selection processes, potentially linked to altered mTEC organization and reduced AIRE expression. The observed changes in SP thymocyte proportions may also reflect, at least in part, a relative shift secondary to the reduction in the DP compartment.

**Figure 3.**
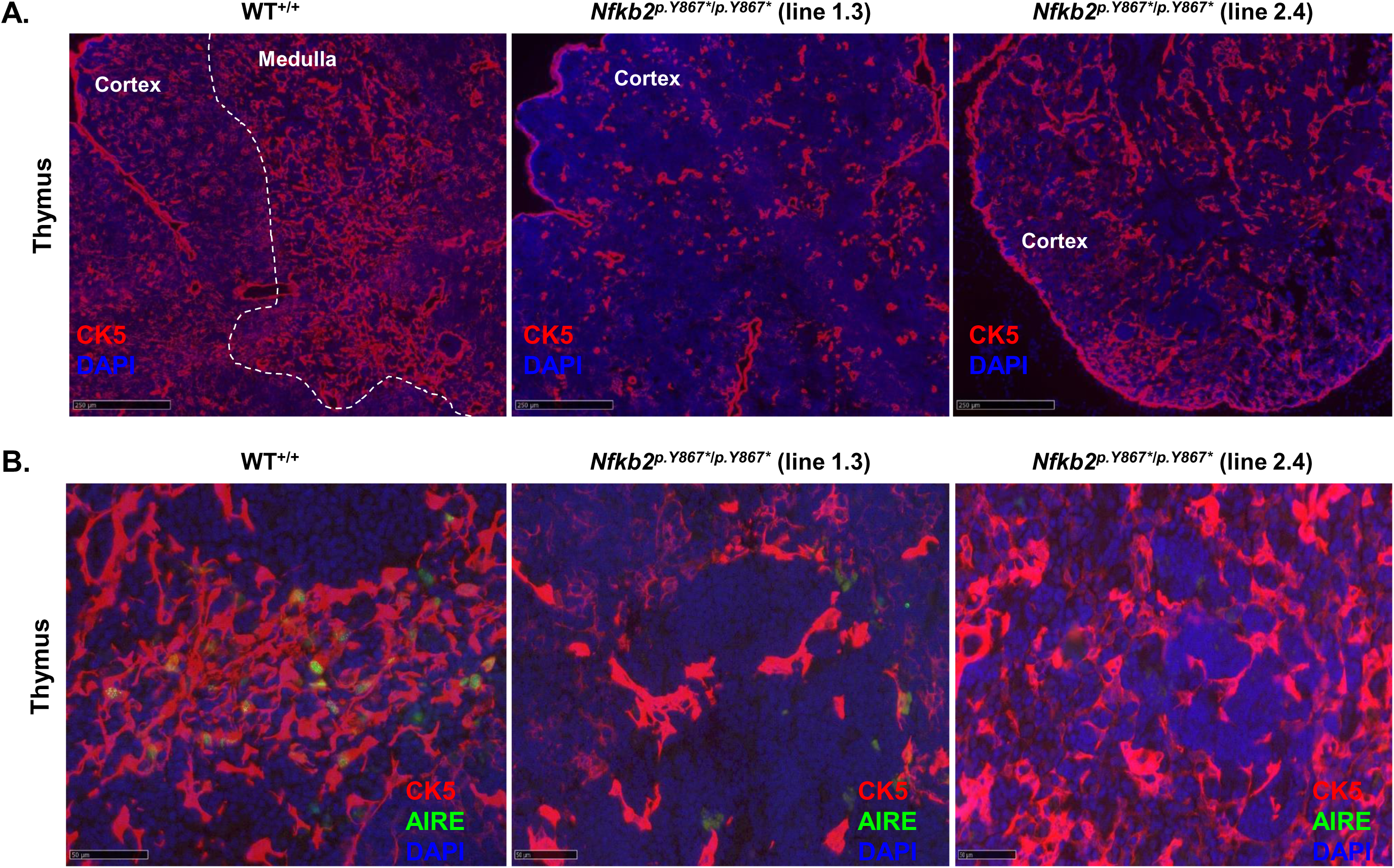
Thymic medullary disorganization and reduced AIRE+ cell compartment in *Nfkb2^p.Y867*/ p.Y867*^*rats. **(A)** Immunofluorescence staining of thymus sections from 3-month-old WT^+/+^ (left panel) and *Nfkb2^p.Y867*/p.Y867*^* rats (line 1.3, middle panel; line 2.4, right panel) showing cytokeratin 5 (CK5, thymic epithelial cells, red) and DAPI (nuclei, blue). The cortical (Cortex) and medullary (Medulla) zones are delineated by a dotted line. Scale bars, 250 µM. **(B)** Immunofluorescence staining of CK5 (red), AIRE (green) and DAPI (nuclei, blue) in thymus sections from 3-month-old WT^+/+^ (left panel) and *Nfkb2^p.Y867*/p.Y867*^*rats (line 1.3, middle panel; line 2.4, right panel). Scale bars, 50 µM.

**Figure 4.**
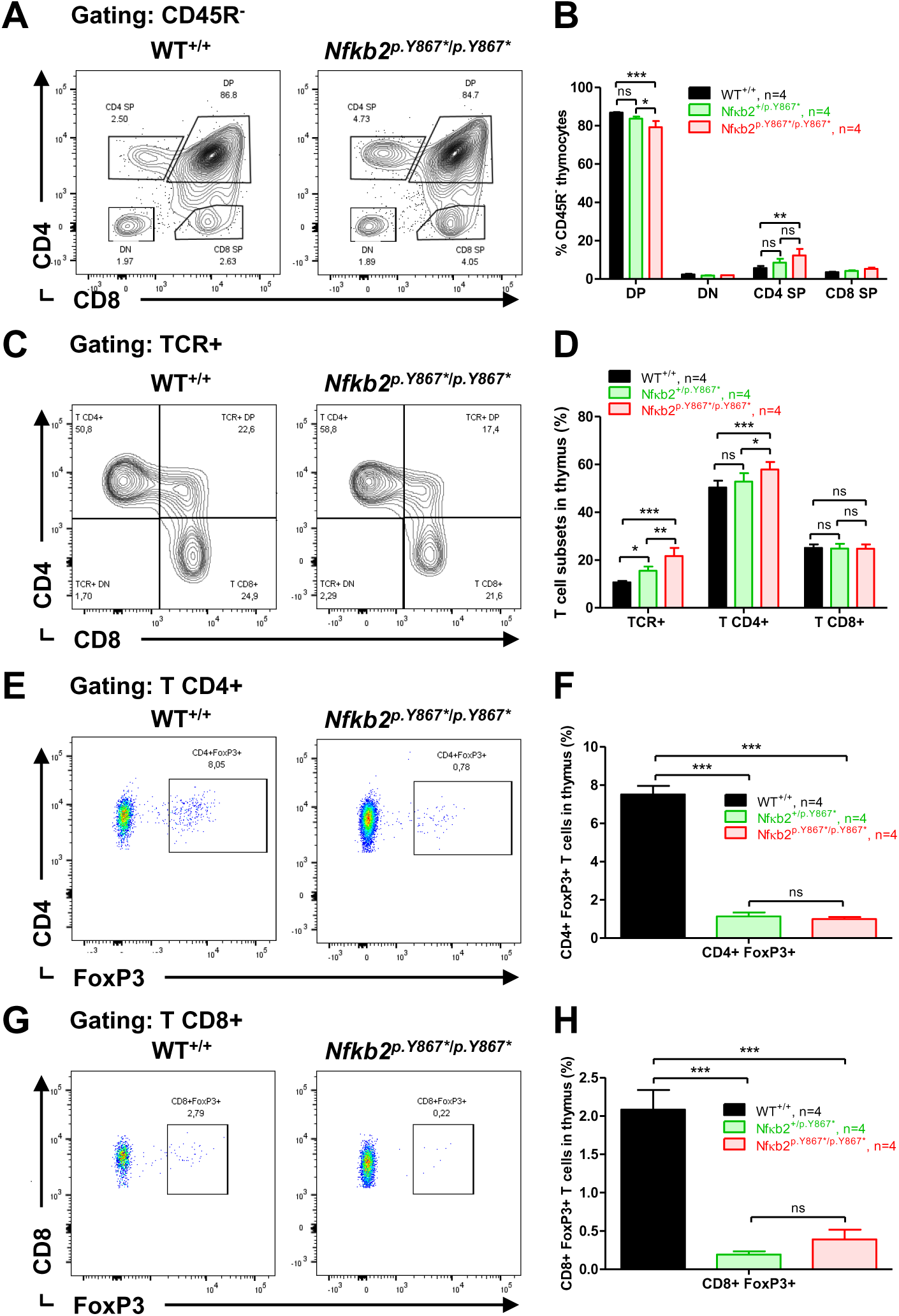
Altered thymocyte development and reduced thymic Treg populations in *Nfkb2^p.Y867*/ p.Y867*^*rats. **(A-B)** Representative dot plots and quantification of major CD45R^−^ thymocytes subsets (CD4 SP, CD8 SP, DP, DN) in WT^+/+^, *Nfkb2^+/p.Y867*^*and *Nfkb2^p.Y867*/p.Y867*^* rats. **(C-H)** Representative dot plots and quantification of thymic T-cell populations, including TCR^+^, CD4^+^ and CD8^+^ subsets, as well as regulatory T cells (CD4^+^FoxP3^+^ and CD8^+^FoxP3^+^) in WT^+/+^, *Nfkb2^+/p.Y867*^*and *Nfkb2^p.Y867*/p.Y867*^*rats. The percentages of each population are indicated on the dot plots. All analyzed rats were from *NFkB2* mutant line 1.3 and were 3- month-old. Statistical comparison used: one-way (F, H) or two-way (B, D) ANOVA with Bonferroni post test; ns, not significant, *p<0.05, **p<0.01, ***p<0.001.

Altogether, these findings indicate that defective p100 processing compromises the thymic medullary microenvironment, and may consequently alter the differentiation and selection of thymocytes, with potential effects on Treg development.

### Nfkb2 mutation disrupts T-cell development and peripheral T-cell homeostasis of *_Nfkb2_p.Y867*/ p.Y867\**_rats_

We next evaluated the impact of thymic alterations on T-cell subsets across central and peripheral compartments.

Phenotypic analyses of T cell populations revealed a significant increase in the frequency of TCR^+^ cells in the thymus and spleen, in both heterozygous and homozygous rats, compared to controls **(Fig. 4D-5B),** as well as in peripheral blood only in homozygous rats (**supplemental Figure 8D)**. These findings suggest a global expansion of the T-cell compartment across the tissues analyzed. Study of T cell subpopulations showed an increase in CD4^+^ T lymphocytes in the thymus in homozygous rats **(Fig. 4C-4D)**. In the spleen, a downward trend in this population was observed, although this did not reach statistical significance **(Fig.5A-5B)**, while no notable changes were detected in peripheral blood **(supplemental Figures 8C-8D)**. These results suggest that the alterations in the CD4^+^ compartment are primarily localized to the thymus. Regarding CD8^+^ lymphocytes, no significant differences were observed between HE, HO and control rats in any of the compartments studied **(Fig. 4C-4D**; **Fig.5A-5B; supplemental Figures 8C-8D)**, although a slight upward trend was observed in spleen of homozygous rats **(Fig.5A-5B)**.

**Figure 5.**
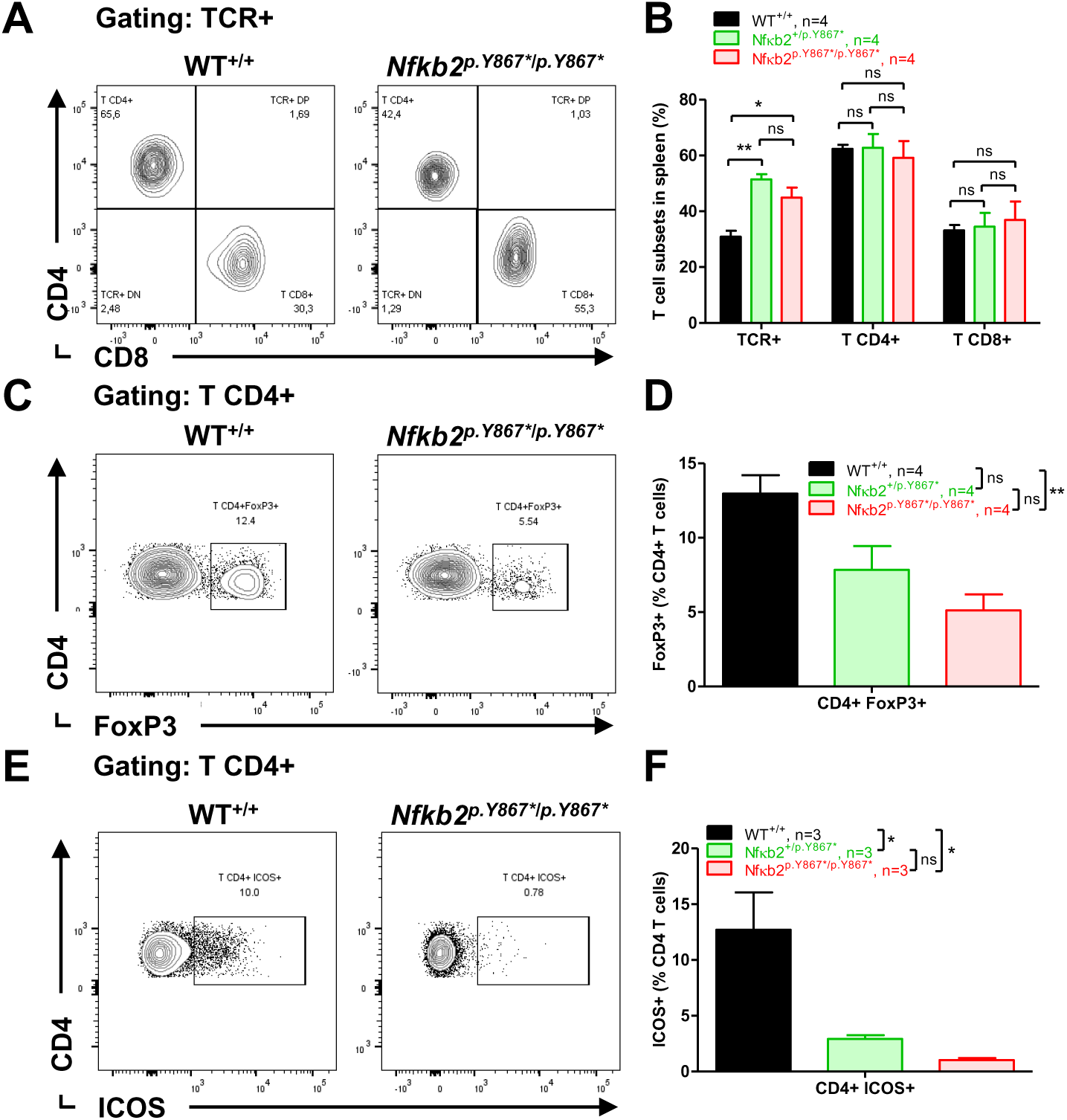
Reduced splenic Treg cells and ICOS expression on CD4+ T cells in *Nfkb2^p.Y867*/ p.Y867*^*rats. **(A-D)** Representative dot plots and quantification of splenic T-cell populations, including TCR^+^, CD4^+^ and CD8^+^ subsets, as well as CD4^+^FoxP3^+^ regulatory T cells and CD4^+^ICOS^+^ T lymphocytes in WT^+/+^, *Nfkb2^+/p.Y867*^*and *Nfkb2^p.Y867*/p.Y867*^* rats. The percentages of each population are indicated on the dot plots. All analyzed rats were from *NFkB2* mutant line 1.3 and were 3-month-old. Statistical comparison used: one-way (D, F) or two-way (B) ANOVA with Bonferroni post-test; ns, not significant, *p<0.05, **p<0.01.

Analysis of regulatory cell populations revealed a marked decrease in Foxp3^+^ regulatory T cells (Tregs) in both HE and HO rats. This decrease was particularly pronounced in the thymus, with an approximately 10-fold reduction in both CD4^+^ Tregs **(Fig.4E-4F)** and CD8^+^ Tregs **(Fig.4G-4H)**. A significant decrease in CD4*^+^* Tregs was also observed in the spleen, more pronounced in homozygous (∼2.5-fold) than in heterozygous animals (∼1.65-fold) **(Fig.5C-5D)**, as well as in the peripheral blood of these same animal groups (∼4-fold) **(supplemental Figures 8E-8F)**. These findings indicate a reduction in Treg cells, consistent with altered immune regulatory homeostasis, particularly in the thymus.

We next examined ICOS expression on peripheral CD4^+^ T cells. Phenotypic analysis of CD4^+^ T cells revealed a significant decrease in ICOS expression in both spleen **(Fig.5E-5F)** and peripheral blood **(supplemental Figures 8G-8H)** of HE and HO rats, as shown by reduced frequencies of ICOS^+^CD4^+^ T cells of approximatively 3.5-fold and 8-fold in the spleen of heterozygous and homozygous rats, respectively **(Fig.5F)**, and of 3-fold and 11.5-fold in the peripheral blood of the same animals **(supplemental Figure 8F)**.

Together, these findings demonstrate a profound alteration of the peripheral CD4^+^ helper T-cell compartment, likely compromising the cellular interactions required for efficient B-cell activation and differentiation. We next examined B-cell development and humoral immune responses in *Nfkb2^p.Y867*^* mutant rats.

### Impairment of peripheral B cell maturation and humoral function in *_Nfkb2_p.Y867*/p.Y867\**_rats_

The marked disorganization of splenic architecture, including the absence of B-cell follicles and marginal zone organization **(Fig. 2C)**, is suggestive of alterations in the splenic microenvironment, which is normally maintained by stromal cells. In the absence of overt splenomegaly **(Supplemental figure 7D),** homozygous rats nevertheless exhibited a 2-fold decrease in total splenic cellularity **(Supplemental figure 7E)**. This reduction appeared to largely spare the CD45^+^ compartment **(Supplemental figure 7F)**, suggesting a relative preservation of hematopoietic leukocytes. In contrast, the decrease in total cellularity may reflect a preferential impact on CD45^−^ populations, including stromal and/or erythroid compartments, in line with the observed disruption of splenic architecture.

We next investigated the impact of these alterations on peripheral B-cell maturation and humoral immune responses in *Nfkb2*-deficient rats. Immunophenotypic analysis further demonstrate a significant reduction in the CD45RA^+^ B-cell population, associated with a pronounced decrease in mature IgD^+^ B cells **(Fig. 6A-6B)**. In contrast, an accumulation of transitional B cells displaying an IgM^+^IgD^−^ phenotype was observed **(Fig. 6A-6B)**, suggesting a developmental blockade at an early stage of B cell maturation. A similar immunophenotypic profile was observed in the blood compartment **(Fig. 6C-6D)**.

**Figure 6.**
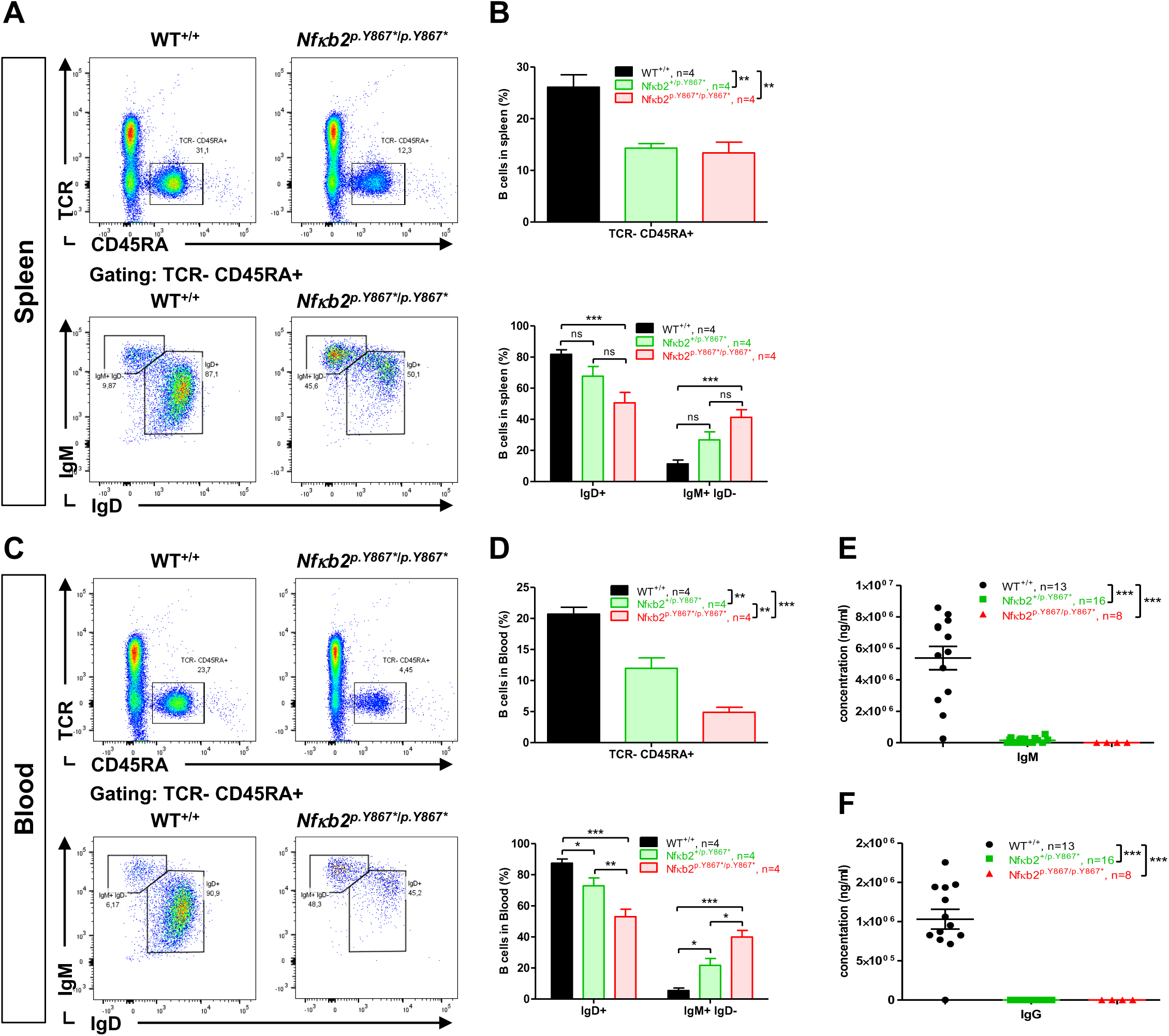
Altered peripheral B-cell subsets and impaired humoral immunity in *Nfkb2^p.Y867*/ p.Y867*^*rats. **(A-D)** Representative dot plots and quantification of TCR^−^CD45RA^+^ B cells (upper dot plots, upper graph), IgM^+^IgD^−^ and IgD^+^ B cells subsets (lower dot plots, lower graph) in spleen (A- B) and peripheral blood (C-D) of 3-month-old WT^+/+^, *Nfkb2^+/p.Y867*^*and *Nfkb2^p.Y867*/p.Y867*^* rats. The percentages of each population are indicated on the dot plots. **(E-F)** Quantitative detection of total IgM and IgG levels in sera of WT^+/+^, *Nfkb2^+/p.Y867*^*and *Nfkb2^p.Y867*/p.Y867*^* rats. Animals were from 2 to 7 months of age. All analyzed rats were from *NFkB2* mutant line 1.3. Statistical comparison used: one-way (B, D, upper graphs; E, F) or two-way (B, D, lower graphs) ANOVA with Bonferroni post-test; ns, not significant, *p<0.05, **p<0.01, ***p<0.001.

Consistent with these cellular alterations, serological analysis showed a complete absence of detectable IgM **(Fig. 6E)** and IgG **(Fig. 6F)** in the serum of *Nfkb2^+/ p.Y867*^*and *Nfkb2^p.Y867*/^ p.Y867\** _rats._

Together, these findings demonstrate a severe impairment of peripheral B-cell maturation and a profound defect in humoral immune responses in rats carrying the homozygous p.Y867* mutation.

## Discussion

Numerous pathogenic variants affecting the C-terminal region of the *NFKB2* gene have been identified in humans and are associated with a heterogeneous spectrum of clinical manifestations, including common variable immunodeficiency (CVID), antibody deficiency, autoimmune manifestations and autoantibodies, as well as endocrine abnormalities such as adrenal insufficiency and pituitary hormone deficiencies [1,5,23–26]. Murine models carrying homologous mutation near phosphorylation sites (e.g *Nfkb2^Lym1^*mice carrying p.Tyr868*- Nfkb2; c.2601T>A) only partially reproduce human phenotypes [1,10,16], as immunodeficiency but not endocrine defects, thus limiting their usefulness for studying associated pathogenic mechanisms. In this regard, the rat is a relevant alternative model, as it has often proved to be more representative of human situations than mice [27]. In this context, we have developed a rat model carrying a c.2598C>A mutation in the *Nfkb2* gene introducing a premature stop codon at Y867 in the C-terminal sequence, and predicting to result in a truncated p100 protein. This mutation is equivalent to the human heterozygous missense variants (c.2602T>A:p.Y868N and c.2603A>G:p.Y868C) identified in three male and one female individuals with CVID [5,28].

Rats carrying the p.Y867* mutation, exhibited impaired maturation of p100 into the active p52 form, resulting in an accumulation of the p100 precursor in CD3-CD28-stimulated splenocytes. These findings are consistent with previous observation in CD40L-stimulated HEK293 cells transfected with the p.Y868N mutant [28] and in *Nfkb2^Lym1^* mice [10,29]. The affected residue is located within the C-terminal region of p100, adjacent to the highly conserved serine residues S866 and S870, which are critical for phosphorylation, ubiquitination and subsequent proteasomal processing of p100 into p52, key steps in activation of the non-canonical NF-κB pathway [30]. Previous studies have shown that mutations affecting S866, S870, or neighboring residues, can impair p100 phosphorylation and processing [23,31–35]. In our study, although the precise molecular consequences of the mutation were not directly investigated, the observed p100 accumulation strongly supports a defect in p100 processing.

RelB expression was not altered in mutant cells. However, RelB accumulation following anti- CD3/anti-CD28 stimulation was observed, raising the possibility of impaired nuclear translocation, as previously suggested in the murine model [29]. Additional studies examining RelB subcellular localization would be required to formally establish this mechanism.

In contrast, the processing of p105 into p50 appeared to be preserved, indicating that the defect is selective for p100 maturation rather than reflecting a global impairment of proteasomal processing within the NF-kB signaling network. Together, these findings support a preferential disruption of the non-canonical NF-kB pathway, notably characterized by the accumulation of immature p100 without evidence of a marked “super-repressor” effect on canonical NF-κB signaling, in contrast to observations reported in mice carrying the homologous mutation [10]. This discrepancy could be due to differences in species, cellular context, and stimulation conditions between the murine studies and our experimental model.

It is well-established that defects in non-canonical NF-κB2 signaling have been associated with a broad spectrum of immune disorders, ranging from severe immunodeficiency to autoimmune and inflammatory diseases, underscoring the critical role of this pathway in maintaining immune homeostasis [36].

Several features of the phenotype observed in our rat model are reminiscent of those described in the *Nfkb2^Lym/Lym1^* mice [10,16]. As observed in the mouse model, the lifespan of homozygous rats was significantly shortened, with a more pronounced phenotype in one of the two lines, in which 50% mortality was observed within the first three months of life. Growth curves were also slightly reduced in homozygous *Nfkb2^p.Y867*^* rats of both sexes, compared with WT littermates. Fertility was not formally assessed in our model, as the lines were maintained by intercrossing heterozygous animals. Nevertheless, we sporadically observed an absence of mature ovarian follicles in young *Nfkb2^p.Y867*/ p.Y867*^* females, which may be consistent with the fertility defects previously reported in the *Nfkb2^Lym/Lym1^* mice The inflammatory liver lesions observed in both heterozygous and homozygous rats increased progressively in severity from HE to HO rats and were associated with a significant reduction in serum albumin levels as early as 3 months of age, which persisted at 7 months. In addition, HO rats exhibited significantly elevated transaminases levels at 7 months, consistent with progressive hepatocellular injury and chronic hepatic inflammation. Together, these findings suggest that the hepatic phenotype develops progressively, with an early alteration in liver function followed by more overt hepatocyte injury. The findings are consistent with previous studies describing hepatic inflammation in the *Nfkb2^Lym1^* model [10].

Profound abnormalities of the lymphoid compartment were also identified, including the absence of mesenteric and peripheral lymph nodes, together with hypoplasia of the spleen and thymus. Furthermore, homozygous rats displayed sever disruption of splenic architecture, characterized by the absence of B follicles and marginal zones, indicating a profound defect in the organization of the secondary lymphoid tissues.

The phenotype observed in our model closely resembles that described in *Nfkb2^Lym1/Lym1^* mice [10] and *Nfkb2*-deficient mice [6,8], which exhibit profound defects in lymphoid organ development and organization, including the absence of peripheral lymph nodes and Peyer’s patches, together with marked abnormalities in thymic and splenic architecture. These phenotypic similarities are consistent with the established role of the non-canonical NF-κB2 pathway in the development and maintenance of primary and secondary lymphoid organs [30,36,37]. In particular, activation of the pathway downstream of LTβR in stromal cells induces the expression of homeostatic chemokines, including CCL19, CCL21 and CXCL13, which contribute to the recruitment and organization of T- and B-cell compartments and to the development and maintenance of secondary lymphoid organs [38].

The structural disorganization of the thymic medulla observed in our model, coupled with a marked reduction in CK5 expression, a well-established marker of medullary epithelial cells, indicates severe alterations within the mTECs compartment and may reflect impaired maintenance, differentiation or survival of these cells. Similar abnormalities have been reported in mice deficient in key components of the non-canonical NF-κB pathway, including RelB, NIK and IKKα. In these mice, the thymic medulla is profoundly disrupted, accompanied by substantial alterations in thymic stromal organization [9,13,15,39].

The concomitant reduction in AIRE expression is particularly relevant in this context, as mature AIRE-expressing mTECs play a pivotal role in the presentation of tissue-restricted antigens and the establishment of central tolerance through negative selection and the generation of thymic regulatory T cells [40].

Previous studies have established a critical link between non-canonical NF-κB signaling and AIRE expression. While *Nfkb2*/p52 deficiency leads to reduced AIRE expression, altered expression of tissue-restricted antigens, impaired central tolerance, and multiorgan autoimmunity [7], the *Lym1* mutation (encoding a truncated and non-processable p100*^Lym1^* protein) results in a near-total loss of *Aire* mRNA associated with fatal thymic medullary hypoplasia [10]. These models collectively demonstrate that both NF-κB2 signaling and p100/p52 homeostasis are required for the establishment of central tolerance through an AIRE-dependent mechanism.

These findings are consistent with earlier studies showing that non-canonical NF-kB signaling is indispensable for the differentiation of mTEC progenitors into mature, functionally competent mTECs capable of supporting negative selection and regulatory T cell development [41,42].

Consistent with the thymic epithelial abnormalities, the reduction in double-positive (DP) thymocytes could reflect impaired thymocyte differentiation or survival, within an altered thymic microenvironment. In contrast, the increased proportion of CD4 single-positive (SP) thymocytes indicates that later stage of thymic selection may be dysregulated. Importantly, this accumulation of CD4-SP cells was accompanied by a reduction in FoxP3^+^ regulatory T cells (Tregs), a population whose development critically depends on signals delivered by mature mTECs within the medulla [43,44], without causing a major defect in conventional thymocytes. This decrease could therefore be linked, at least in part, to alterations in the mTEC/AIRE compartment and changes in thymic selection. However, given the complex regulation of Treg development, which involves both thymic epithelial cells and hematopietic cells, our data do not establish a direct causal link between reduced AIRE expression and the Treg deficiency.

Overall, the combined alterations involving CK5, AIRE, DP and CD4 SP thymocytes, and Tregs indicate a global disruption of thymic homeostasis associated with altered NFkB2 signaling.

Like the reduction observed in the thymus, FoxP3^+^ Treg cells were also markedly reduced in peripheral compartments, including spleen and peripheral blood, despite the relatively preserved development of conventional T cells. The concomitant decrease in ICOS- expressing CD4^+^ T cells suggests a broader alteration of peripheral CD4^+^ T-cell homeostasis in *Nfkb2-*deficient rats. These findings are consistent with the broader alterations in T-cell subsets reported in patients with pathogenic *NFKB2* variants, while also highlighting interspecies differences regarding the impact of Nfkb2 deficiency on T-cell homeostasis [5]

In contrast, *Nfkb2-*deficient mice exhibited preserved Treg numbers in thymus and even increased frequencies in the periphery, accompanied by enhanced in vivo activation and proliferation [45]. Together with the cell-intrinsic role of NIK, these findings suggest that non-canonical NF-kB signaling contributes to Treg development and peripheral homeostasis. Consistent with this, NIK deficiency intrinsically impairs Treg generation and compromises immune tolerance [46]. Interestingly, the *Nfkb2^Lym1^*phenotype appears more complex. Wirasinha et al. showed that, despite impaired Treg generation, *Nfkb2^+/Lym1^* Tregs retain substantial suppressive capacity in vivo, although they are less effective at controlling the spontaneous activation of CD4^+^ T cells [16]. Moreover, the autoimmune phenotype was strongly influenced by abnormalities in the non-hematopoietic compartment, highlighting the contribution of the thymic environment to this alteration [16].

These findings indicate that alterations in Nfkb2 can lead to markedly divergent effects on peripheral CD4^+^ T-cell homeostasis, ranging from severe Treg depletion as observed in our model and discussed by Klemann et al. [5], to T-cell hyperactivation as reported by Dhar et al. [45].

The alterations observed within the B-cell compartment are also consistent with defective non-canonical NF-kB signaling. This pathway plays a crucial role in B-cell survival, maturation and functional responses, particularly downstream of BAFF-R and CD40 [36]. Mouse models with genetic deletions of p52 [8], RelB [47], or NIK [48,49] consistently exhibited splenic microenvironment disorganization and an arrest in B-cell differentiation. Like the *Nfkb2^Lym1^* strain [10], our model reveals a maturation defect in splenic and peripheral B-cell compartments, functionally resulting in a collapse of the humoral responses characterized by the absence of circulating IgM and IgG. The lack of detectable autoantibodies may be explained, at least in part, by the profound hypogammaglobulinemia observed in these rats.

Importantly, this molecular and functional phenotype directly recapitulates human immunodeficiencies caused by heterozygous pathogenic variants in the C-terminal region of NFKB2; these variants disrupt p100 maturation and lead to DAVID syndrome or CVID [5,50]. Taken together, these data support the concept that the regulated maturation of p100 is a key, evolutionarily conserved element for ensuring functional humoral competence.

Interestingly, despite the severe immune phenotype observed in our *Nfkb2^p.Y867*^*rat model, we did not identify any significant abnormalities in pituitary development or hormone production. These findings are consistent with previous studies on *Nfkb2* mouse models, notably the *Nfkb2^Lym1^* strain, which failed to reproduce the pituitary abnormalities characteristic of patients with NF-κB2-associated DAVID syndrome [1,51]. This apparent discrepancy highlights a significant interspecies difference regarding the role of NF-κB2 signaling in pituitary development and function. The absence of a pituitary phenotype in mouse and rat models, despite the well-established endocrine manifestations in patients, suggests that the mechanisms underlying the endocrine component of DAVID syndrome may not be fully conserved between rodents and humans. Recent studies using human pituitary organoids carrying NF-κB2 mutations provide evidence of a direct role of NF-κB2 in human pituitary differentiation, thereby reinforcing this hypothesis [51].

Collectively, our findings demonstrate that the pathogenic p.Y867*-Nfkb2 variant induces a systemic combined immunodeficiency phenotype in rats, deeply altering multiple processes regulated by the non-canonical NF-kB pathway. Mechanistically, this alteration in NF-kB2 signaling results from a defect in p100 protein maturation, exacerbated by the accumulation of the precursor. Our study establishes this new model as a valuable tool for deciphering the crucial role of the non-canonical NF-kB signaling in the organization of immune system and in adaptive immunity.

## Methods

### Animals

Sprague-Dawley (SD/Crl) rats were the only strain used and were sourced from JanvierLabs (Le Genest-Saint-Isle, France). All the animal care and procedures performed in this study were approved by the Animal Experimentation Ethics Committee of the Pays de la Loire region, France, in accordance with the guidelines from the French National Research Council for the Care and Use of Laboratory Animals (Permit Numbers: CEEA-PdL-2015-692; APAFIS#54911).

### spCas9, sgRNA preparation and ssODN

sgRNA was designed to target the following sequences: TGCCTACGGGAGCCAGTCAG*TGG.* crRNA was purchased from IDT and hybridized with tracrRNA to form sgRNA, then the ribonucleoprotein complex was done with sp Hifi Cas9 V3 from IDT.

The ssODN-p.Y867*-EcoRI (100 nt), was purchased from IDT in dry form, dissolved in nuclease free-water at 1mM and stored at -20°C until use.

ssODN-p.Y867*-EcoRI sequence: 5’ggcttgactcccccctgtcccctggcagaggtgaaagaagacagtgcTtGA**gaattc**caaagcgtcgagcaggaggcagaga agctgtgtccaccccctgagcctccaggagggctctgc-3’

### Collect and electroporation of rat one-cell embryos

Prepubescent females (4-5 weeks old) were super-ovulated with pregnant mare serum gonadotropin (30 IU; Intervet, France) and followed 48 hours later with human chorionic gonadotropin (20 IU; Intervet, France) before breeding. Fertilized 1-cell stage embryos were collected and kept in M16 medium at 37 °C under 5% CO2 until electroporation.

sgRNA and Cas9 protein were incubated at room temperature for 10 min to allow formation of ribonucleoprotein complexes. ssODN was added to sgRNA/Cas9 protein complexes. The mixture was kept on ice until electroporation. Electroporation was performed using the NEPA21 system (NEPA GENE Co. Ltd, Sonidel, Ireland) as previously described [21]. Electroporated embryos were maintained under 5% CO_2_ at 37 °C until implantation. Surviving embryos were implanted the same day in the oviduct of pseudo-pregnant females (0.5 dpc) and allowed to develop until birth.

### Genotyping

Briefly, ear biopsies from 8- to 10-day-old rats were digested overnight at 56 °C, in 500 µL of tissue digestion buffer (0.1 M Tris-HCl pH 8.3, 5 mM EDTA, 0.2% SDS, 0.2 M NaCl, 100 µg/mL proteinase K). The CRISPRs nuclease targeted regions were PCR amplified from diluted lysed samples (1/20) with a high-fidelity polymerase (Herculase II fusion polymerase). To detect gene edition, locus specific primers *Nfkb2* (r*Nfkb2*For2: gtctattggaagccttgtccga / r*Nfkb2*Rev2: tgccctttgtatagccttcacc) are used and mutations were analyzed by HMA-CE [52] and direct sequencing of PCR products. ssODN insertion introduces a premature termination codon and a EcoRI restriction site in the PCR product, which allows easy detection of KI animals by simple digestion.

### In vitro activation of splenocytes

For western blot analyses, splenocytes isolated from *Nfkb2^p.Y867*/ p.Y867*^*, *Nfkb2^+/ p.Y867*^* and WT littermate rats were stimulated with plate-bound anti-CD3 (1 µg/ml; clone G4.18) plus anti- CD28 (1 µg/ml; clone JJ319) for 16 h at 37 °C under 5% CO2. Cells were pelleted for 7 min at 1400 rpm at 4 °C and washed twice with cold PBS. Cell pellets were then resuspended in RIPA buffer (100 µl for 10 × 10^6^ cells) containing protease inhibitor cocktail and incubated for 30 min on ice. After centrifugation for 10 min at 10,000 rpm at 4 °C, supernatant was harvested and protein concentration estimated using a Pierce™ BCA Protein Assay Kit (#23225, ThermoFisher Scientific).

### Western Blot

Ten micrograms of total protein were run on a 4-15% Mini PROTEAN TGX precast gels and then transferred onto a nitrocellulose membrane using the Trans-Blot Turbo Transfer System (Bio-Rad Laboratories). Membranes were blocked for 2 h at room temperature on constant agitation with 5% BSA in a Tris-buffered saline with 0.1% Tween-20 (TBST) solution.

Membranes were then incubated overnight at 4 °C on constant agitation with rabbit anti-p52 (#4882; Cell Signaling Technology), anti-p50 (#13586S; Cell Signaling Technology), anti-RelB (#10544S; Cell Signaling Technology), anti-RelA (#8242S; Cell Signaling Technology) or mouse anti-β actin antibody diluted 1 : 1000 in 1% BSA in TBST solution. After washing three times with TBST solution, membranes were incubated for 1 h at room temperature on constant agitation in HRP-goat anti rabbit IgG (H+L) (111-035-144; Jackson Immunoresearch) or HRP-goat anti mouse IgG (H+L) (115-035-146; Jackson Immunoresearch) diluted 1 : 10000 in 1% BSA in TBST solution. Pierce™ ECL Western Blotting substrate (ThermoFisher Scientific) was used as a substrate and chemiluminescence detected on a FujiFilm LAS-4000 camera.

### Flow cytometry

T cell subsets, B cells subsets in single-cell suspensions isolated from spleen, thymus and blood were analyzed by flow cytometry. Antibodies and reagents used for immunophenotyping are listed in S1 Table. For intracellular detection of Foxp3, cells were stained using Foxp3 staining buffer set (eBioscience) and according manufacturer’s instructions and Foxp3-specific antibody (FJK-16S, Invitrogen. Data were acquired with CantoII flow cytometer (BD) and analyzed with FlowJo software (FlowJo).

### Histological analyses

Organs from *Nfkb2^p.Y867*/ p.Y867*^*, *Nfkb2^+/ p.Y867*^* and WT littermate rats were collected in 4% paraformaldehyde. After 24-48 h of fixation at 4 °C, samples were dehydrated and transferred to 70% ethanol for storage if necessary, before being embedded in paraffin, sectioned at 5 μm, and stained with hematoxylin-phloxin-saffron. Images were captured using the NanoZoomer slide scanner (Hamamatsu) and analyzed with NDP.view2 software (Hamamatsu). Lymphocyte infiltration and pathological changes in the tissues were assessed by a pathologist in a blinded manner.

### Immunofluorescence

Seven-micrometer-thick cryosections of *Nfkb2^p.Y867*/ p.Y867*^*, *Nfkb2^+/ p.Y867*^* and WT rat thymus were fixed in 4% paraformaldehyde for 15 min, washed 3 times with PBS, permeabilized in 0.3% Triton X100 for 15 min, and blocked with 10% normal goat serum (NGS) diluted in PBS with 0.1% Triton X-100 and 1% BSA (PBT) for 1 h at room temperature. Sections were then overnight incubated at 4 °C with primary antibodies diluted in 1% NGS in PBT buffer, including anti-cytokeratin 5 (CK5; clone 3P3T9; Biotechne) and anti-AIRE-AF488 (clone 5H12; 53-5934-82; Invitrogen). Sections were washed 3 times with PBS, and then incubated with an anti-rabbit A568 (CK5) diluted in 1% NGS in PBT buffer for 1 h at room temperature. Finally, sections were washed, stained with 1 μg/mL of DAPI for 15 minutes and then mounted with ProLong Gold Antifade reagent (Molecular Probes). Images were collected using NanoZoomer Slide scanner system (Hamamatsu) and analyzed with NDP.view2 software (Hamamatsu).

### Hematological, biochemical, and hormonal parameters

Blood samples were collected and divided into separate tubes. EDTA tubes were used for hematological analyses. Heparin tubes were used for biochemical or hormonal analyses; samples were centifugated for 15 min and sera were then analyzed.

The simultaneous quantification of ACTH, TSH, GH, FSH, LH and prolactin hormones in animals sera (one-third diluted) was performed with a MILLIPLEX kit (RPTMAG-86K-06; Merck) according to the manufacturer’s instructions with a Luminex xMAP detection method.

### Quantitative determination of serum immunoglobulins

Plates were coated overnight at 4 °C with 5 μg/mL of goat polyclonal anti-rat IgM or IgG antibodies (supplied by Jackson ImmunoResearch Laboratories). After several washing steps, saturation with PBS/5% BSA was performed for 2 hours at room temperature. Diluted samples and isotype control (rat IgM, rat IgG supplied by Jackson ImmunoResearch Laboratories), used to determine the standard curves, were incubated for 90 minutes at 37 °C. HRP-conjugated anti-rat IgM or rat IgG (supplied by Jackson ImmunoResearch Laboratories) was incubated for 90 minutes at 37 °C to allow specific detection of rat immunoglobulin isotypes. Tetramethylbenzidine (TMB; BD Biosciences) was used as HRP chromogenic substrate before the reaction was stopped by addition of H2SO4. Optical density at 450 nm was determined using a Tecan Spark plate reader.

### Autoantibodies

Serum autoantibodies against mouse IFNα2 were measured by radioligand binding assay (RLBA) as previously described [53]. Full-length Myc-tagged mouse IFNα2 was synthesized from an expression plasmid (OriGene, MR225508) by *in vitro* transcription and translation using the TNT® Quick Coupled Transcription/Translation System (Promega) in the presence of [^35S]methionine to generate radiolabeled protein. Radiolabeled IFNα2 was incubated with individual serum samples, and antibody-bound antigen was immunoprecipitated using protein A/G Sepharose beads (Cytivia, 17-1279-02/17-0618-02). Following washing, retained radioactivity was quantified by liquid scintillation counting. A commercial anti-Myc antibody (Cell Signaling Technology, 2272) was included as a positive control, and reactions lacking serum were used to determine background signal. Antibody index (AI) was calculated as (sample value − mean blank value)/(positive-control antibody value − mean blank value), as previously described [53]. Samples with an AI greater than three standard deviations above the mean of wild type control sera were considered positive.

For detection of autoantibodies, slides of tissues (pancreas, kidney, liver, colon, and salivary glands) from IgM-deficient rats (to avoid background) were saturated with PBS plus 5% BSA for 30 minutes at room temperature. Sera from *Nfkb2^p.Y867*/ p.Y867*^*, *Nfkb2^+/ p.Y867*^* and WT littermate rats were diluted at 1:5 in PBS/1% BSA and added on the slices. After 30 minutes of incubation at room temperature, slices were washed 3 times with filtered PBS. Autoantibodies were revealed using an FITC–coupled donkey anti-rat IgG antibody (heavy and light chain; Invitrogen). After 30 minutes of incubation with the secondary antibody and another washing step, slices were stained with 2 μg/mL of DAPI for 15 minutes and then mounted with ProLong Gold Antifade reagent (Molecular Probes). Fluorescence was observed using the Nanozoomer slide scanner (Hamamatsu).

## Ethics approval

All the animal care and procedures performed in this study were approved by the Animal Experimentation Ethics Committee of the Pays de la Loire region, France, in accordance with the guidelines from the French National Research Council for the Care and Use of Laboratory Animals (Permit Numbers: CEEA-PdL-2015-692; APAFIS#54911).

## Availability of data and materials

The datasets generated and/or analyzed during the current study are available from the corresponding author on reasonable request.

All data generated or analyzed during this study are included in this published article [and its supplementary information files].

## Authors’ contributions

Conceived and designed the experiments, IA, MG, SR, MSA

Performed the experiments, SR, LT, CU, MM, AY, YP, DY

Analyzed the data, SR, LT, YP, DY

Wrote the paper, SR, IA

Funding acquisition, SR, IA, MG, MSA

All authors reviewed and accepted the submitted version

## Acknowledgments

We thank the local academic laboratories and platforms: GenoA for sequencing experiments, MicroPICell for histological analyses, T. Larcher (APEX) for his expertise in anatomopathology, C. Braudeau (CIMNA) for MILLIPLEX experiments, L2R-CHU Nantes for hematological and biochemical analyses, as well as J. Veziers (SC3M) and F. Blanchard (RMeS) for bone tissues analyses.

