## Supplemental Figures for "A PATHOGENIC NFKB2 VARIANT CAUSES COMBINED IMMUNODEFICIENCY WITH THYMIC AND PERIPHERAL LYMPHOID DEFECTS IN RATS"

### Supplemental Figure 1

A

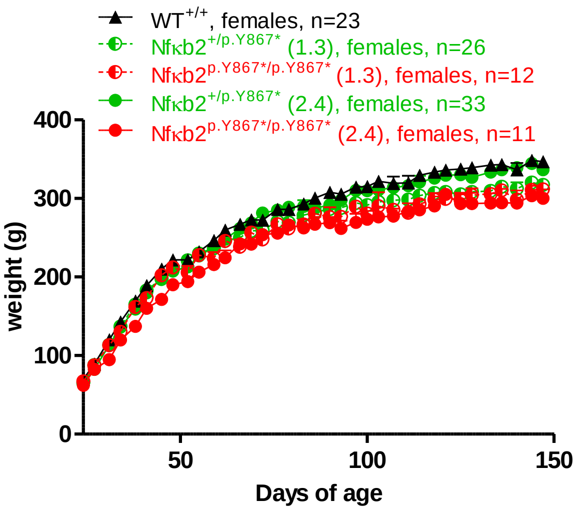


B

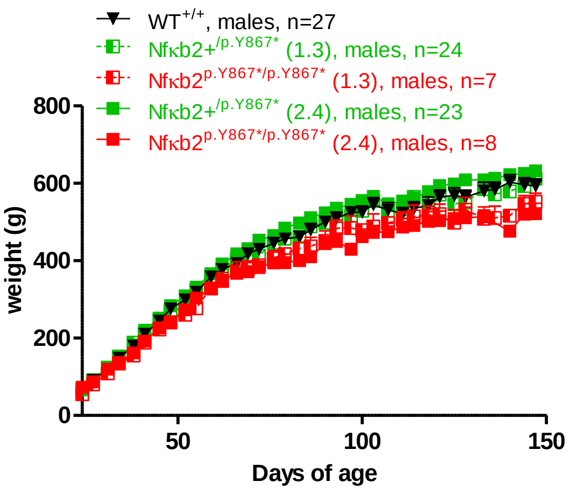


**Comparison of the growth curves of rats of each genotype with those of wild-type groups**
**(A)** The growth curves of *Nfκb2*$^{+/p.Y867*}$ (line 1.3, n=26) (line 2.4, n=33), *Nfκb2*$^{p.Y867*/p.Y867*}$ (line 1.3, n=12) (line 2.4, n=11) and WT$^{+/+}$ (n=23) females.
**(B)** The growth curves of *Nfκb2*$^{+/p.Y867*}$ (line 1.3, n=24) (line 2.4, n=23), *Nfκb2*$^{p.Y867*/p.Y867*}$ (line 1.3, n=7) (line 2.4, n=8) and WT$^{+/+}$ (n=27) males.
Weight was measured two times a week from weaning until 150 days of age. There were not statistical differences.

#### Supplemental Figure 2

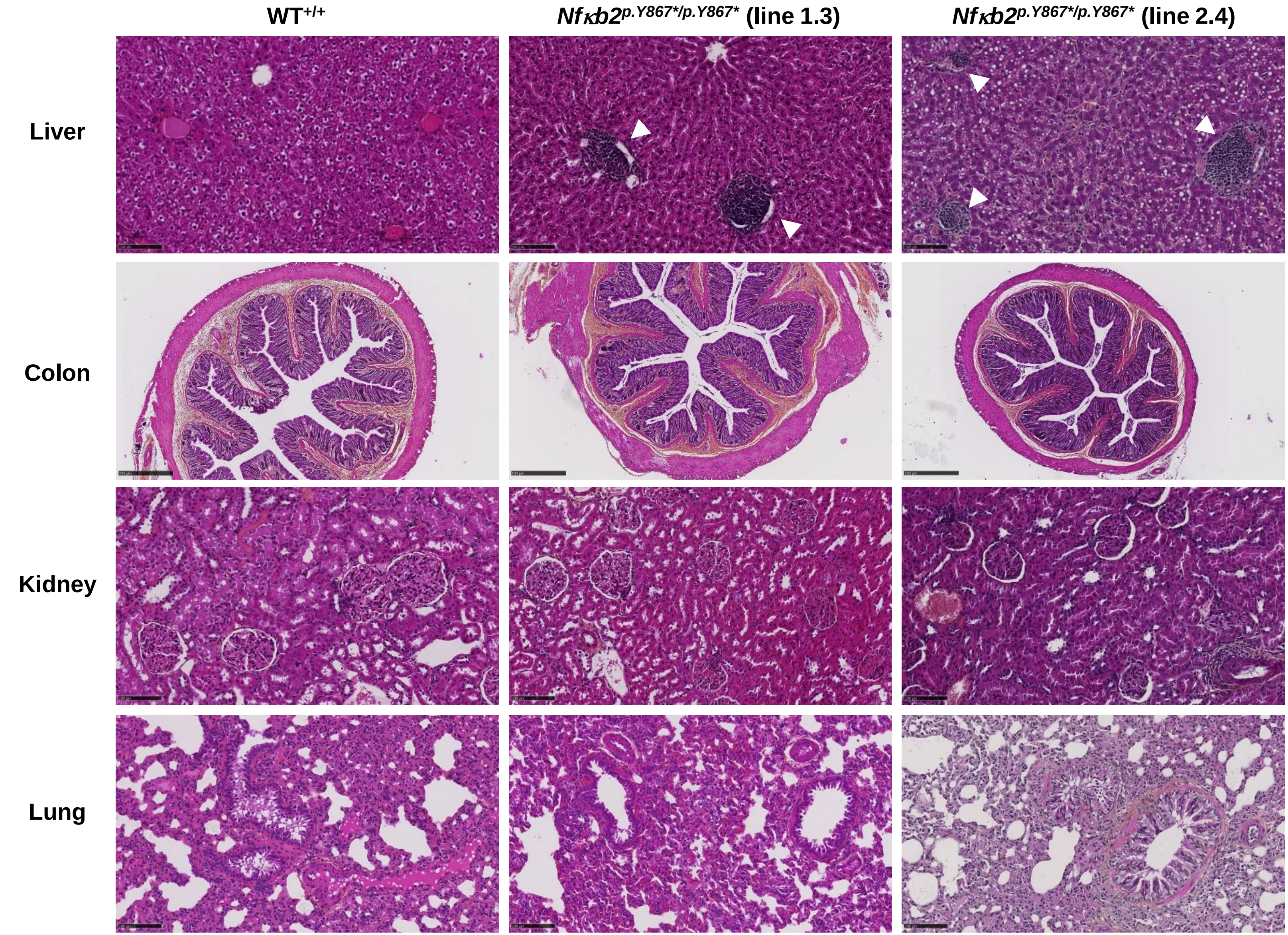


**Histological comparison of tissues between mutated and wild-type rats.** Representative histology of pathological (liver) and normal (colon, kidney, lung) tissues from 3-month-old WT$^{+/+}$ (left panels) and *Nfκb2*$^{p.Y867*/p.Y867*}$ rats (middle panels, line 1.3; right panels, line 2.4). Hematoxylin- phloxine-saffron staining. Inflammatory foci in liver are indicated by white arrows. Scale Bars 100µm (liver) and 500µm (colon, kidney, lung).

### Supplemental Figure 3

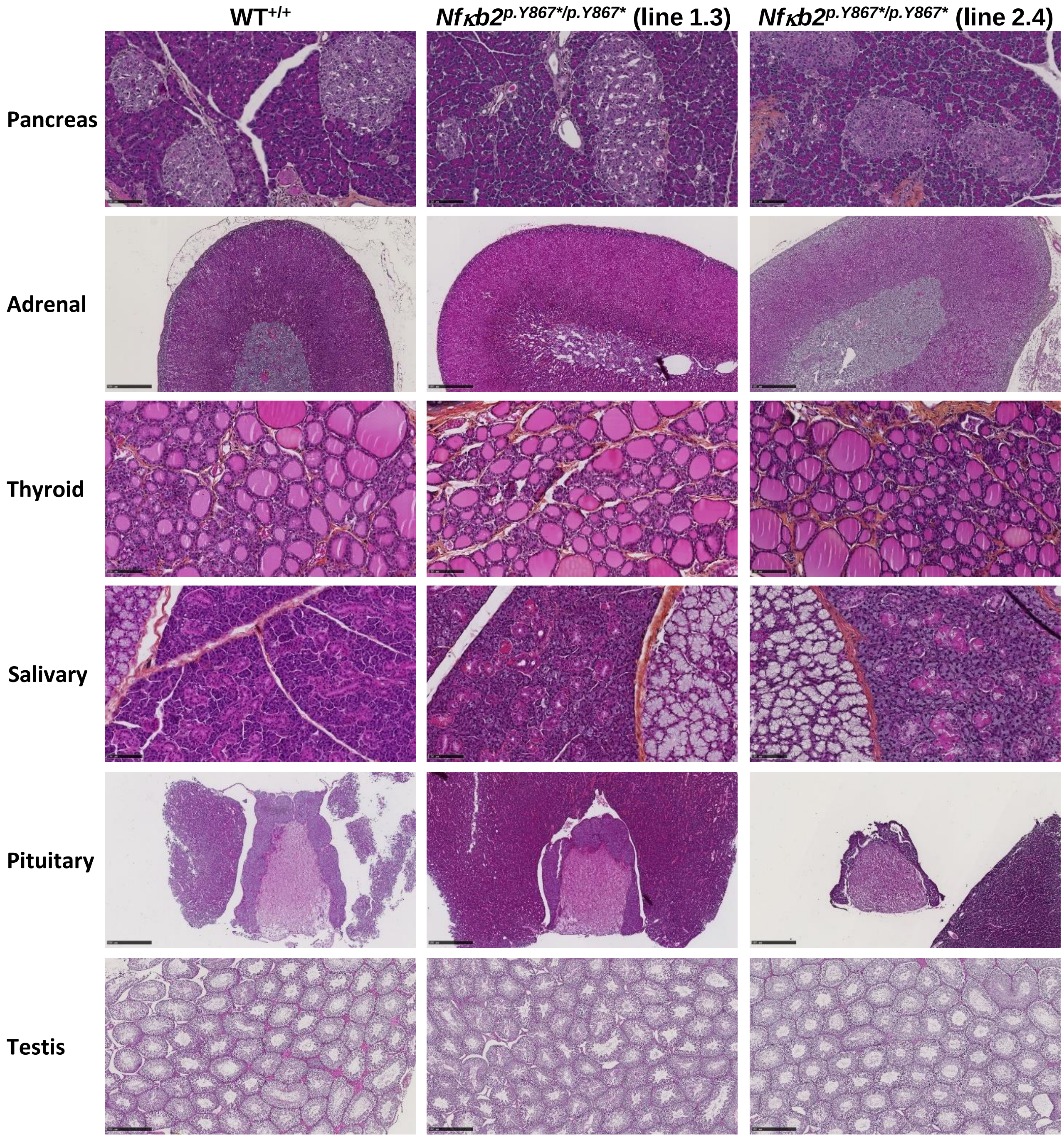


**Histological comparison of endocrine tissues between mutated and wild-type rats.** Representative histology of normal tissue (pancreas, adrenal, thyroid, salivary, pituitary, testis) from 3-month-old WT$^{+/+}$ (left panels) and *Nfκb2*$^{p.Y867*/p.Y867*}$ rats (middle panels, line 1.3; right panels, line 2.4). Hematoxylin-phloxine-saffron staining. Scale Bar 500µm

### Supplemental Figure 4

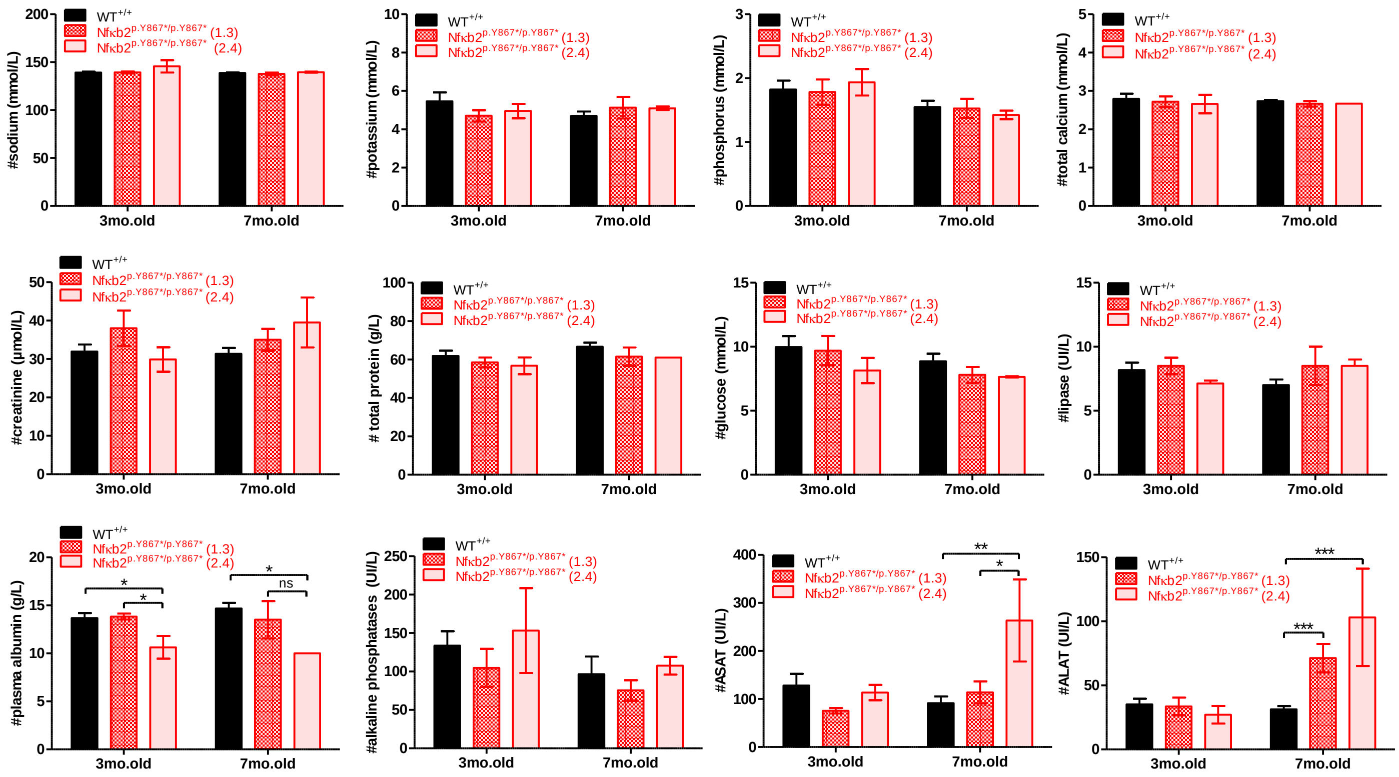


**Analysis of biochemical parameters between mutant and wild-type rats.** Comparison of sodium, potassium, phosphorus, total calcium, creatinine, total protein, glucose, lipase, plasma albumin, alkaline phosphatases and transaminases (ASAT/ALAT) in *Nfκb2*$^{p.Y867*/p.Y867*}$ rats (lines 1.3 and 2.4) with WT$^{+/+}$ rats . Sera were collected at 3 and 7 months of age.

Three-month-old rats: WT$^{+/+}$, n=15; *Nfκb2*$^{p.Y867*/p.Y867*}$ (line 1.3, n=6; line 2.4, n=8).

Seven-month-old rats rats: WT$^{+/+}$, n=15; *Nfκb2*$^{p.Y867*/p.Y867*}$ (line 1.3, n=4; line 2.4, n=2).

Two-way ANOVA with Bonferroni post-test; *p<0.05, **p<0.01, ***p<0.001

### Supplemental Figure 5

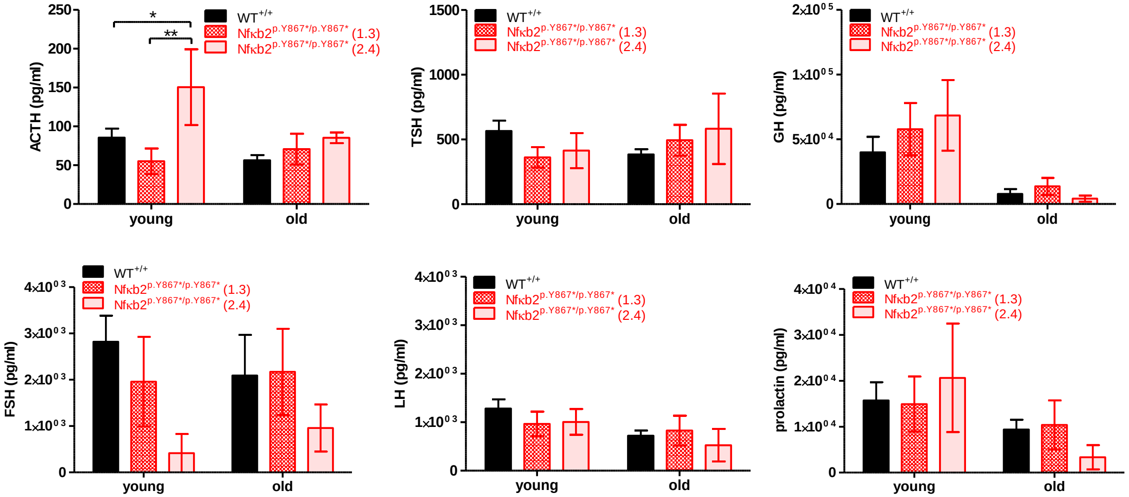


**Comparative analysis of serum pituitary hormones.**

Comparison of ACTH, TSH, GH, FSH, LH and prolactin in *Nfκb2*$^{p.Y867*/p.Y867*}$ rats (lines 1.3 and 2.4) with WT$^{+/+}$ rats . Sera were collected at 3 months (young) and 7-12 months (old) of age (undifferentiated sex).

Young rats: WT$^{+/+}$, n=20; *Nfκb2*$^{p.Y867*/p.Y867*}$ (line 1.3, n=9; line 2.4, n=6).

Old rats: WT$^{+/+}$, n=26; *Nfκb2*$^{p.Y867*/p.Y867*}$ (line 1.3, n=5; line 2.4, n=2).

Two-way ANOVA with Bonferroni post-test; $*p<0.05$, $**p<0.01$

### Supplemental Figure 6

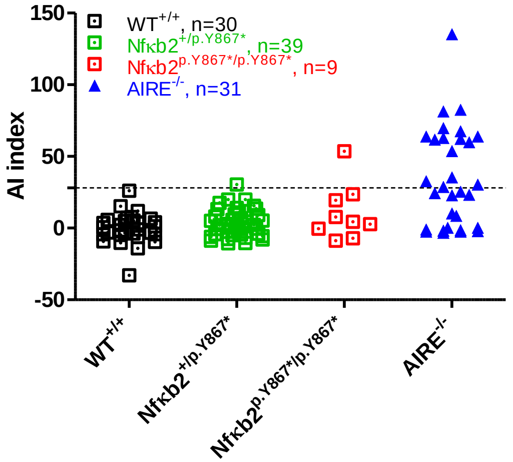


**Analysis of Anti-IFNa2 auto-antibodies.**

Anti-IFNa2 auto-antibodies detection by Radioligand binding assay (RLBA) in sera from aged (7- to 14-month-old rats) *Nfkb2*$^{+/p.Y867*}$ (n=39), *Nfkb2*$^{p.Y867*/p.Y867*}$ (n=9) (both lines combined) and WT$^{+/+}$rats (n=30). AIRE$^{-/-}$ sera were used as positive control (n=31) (7- to 23-month-old rats). AI = Autoantibody Index; AI Cutoff (28.08) was calculated as average of healthy controls + 3 SEM; Antibody Control is anti-Myc antibody

### Supplemental Figure 7

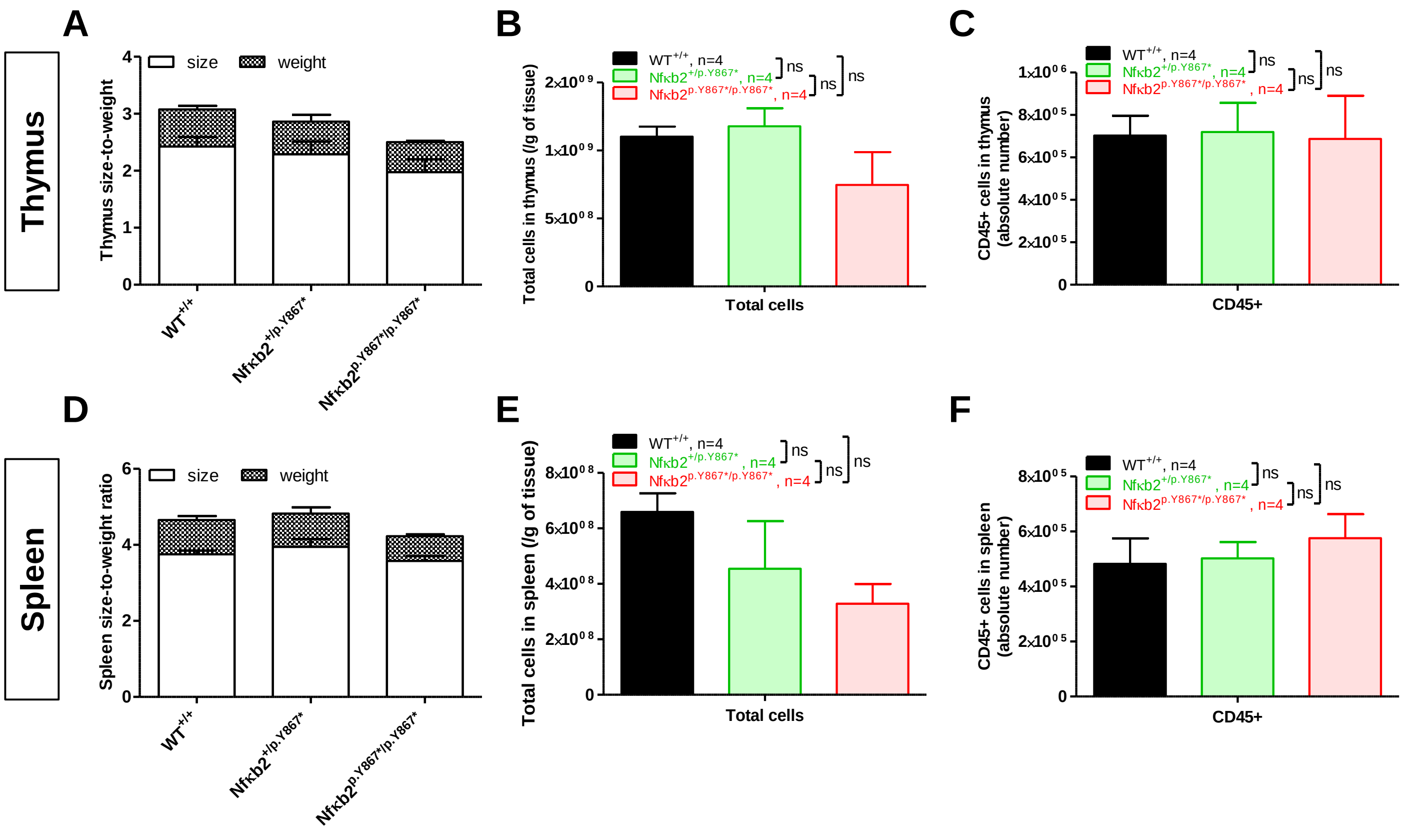


**Morphological and cellular comparison of the spleen and thymus between mutant and wild-type rats.**
Comparison of thymus **(A-C)** and spleen **(D-F)** in *Nfκb2*$^{+/p.Y867*}$ and *Nfκb2*$^{p.Y867*/p.Y867*}$ rats (line 1.3) versus WT$^{+/+}$ rats.
**(A, D)** Organ size-to-weight ratio.
**(B, E)** Total cell number per gram of tissue
**(C, F)** CD45$^{+}$ cell counts (absolute number)
Data pooled from 3-month-old rats (N=4) and compared using one-way ANOVA; ns, not significant

### Supplemental Figure 8

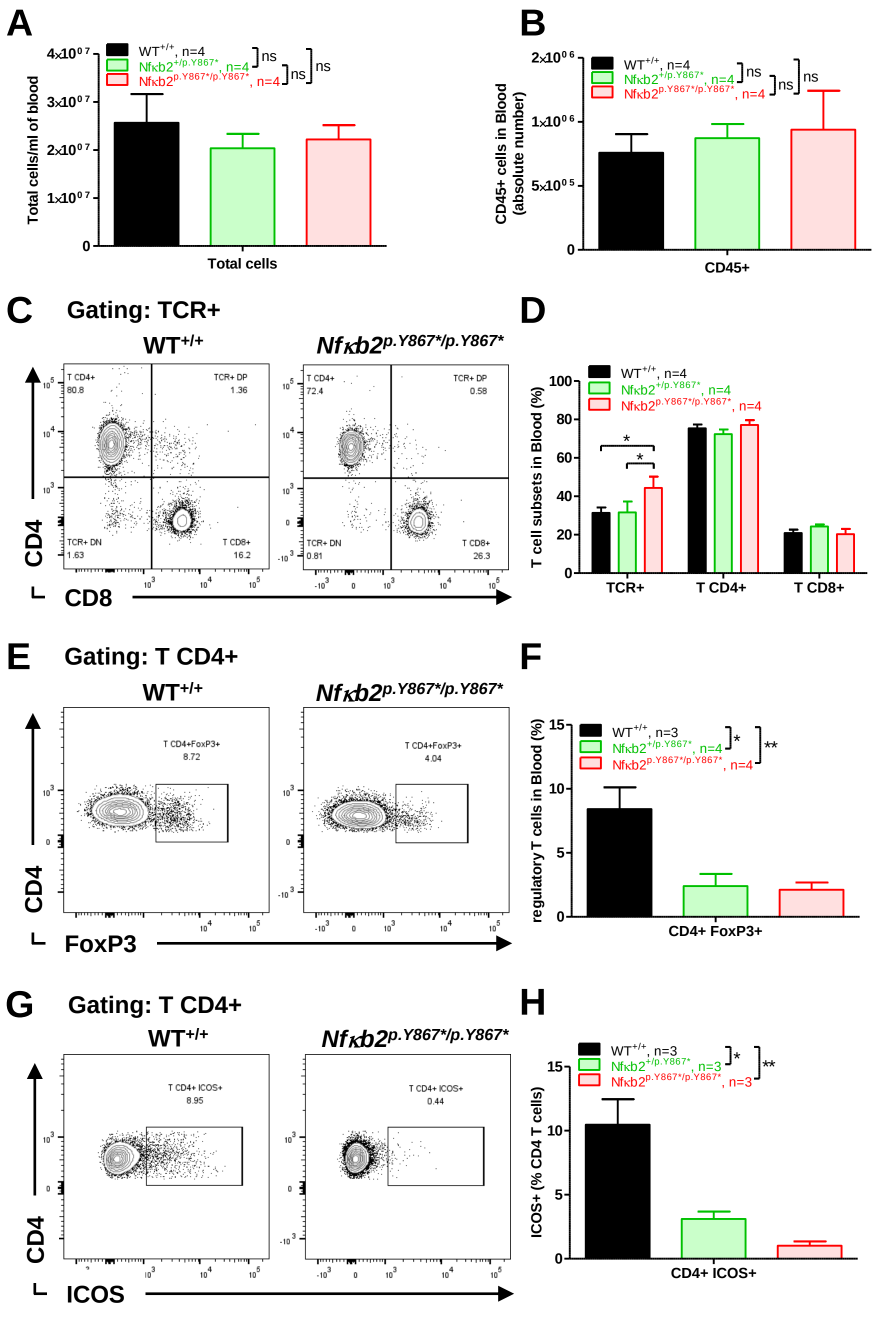


**Altered peripheral T-cell phenotype in *Nfkb2*$^{p.Y867*/p.Y867*}$ rats**

**(A, B)** Total peripheral blood cell count and absolute number of $CD45^+$ cells

**(C, D)** Representative plots and quantification of $TCR^+$, $CD4^+$ and $CD8^+$ T cells

**(E, F)** Representative plots and quantification of $CD4^+FoxP3^+$ Treg cells

**(G, H)** Representative plots and quantification of $ICOS^+$ $CD4^+$ T cells

Data pooled from 3-month-old rats and compared using one-way ANOVA (A, B, F, H) or two-way ANOVA (D), with Bonferroni post-test (D, F, H); ns, not significant, *$p<0.05$, **$p<0.01$.
