## Supplemental Table 1 for "A PATHOGENIC NFKB2 VARIANT CAUSES COMBINED IMMUNODEFICIENCY WITH THYMIC AND PERIPHERAL LYMPHOID DEFECTS IN RATS"

**S1 Table. List of antibodies and reagents used for flow cytometry**

Target Antigen, clones, fluorochromes, manufacturers, catalog numbers, Research resource Identifiers (RRID), and working dilutions used for immunophenotyping.

N/A (Not Applicable)

| **Target**  **Antigen** | **Clone** | **Fluorochrome** | **Manufacturer** | **Catalog No.** | **RRID** | **Dilution** |
| --- | --- | --- | --- | --- | --- | --- |
| CD45 | OX1 | APC-Cy7 | BD Biosciences | 561586 | AB_10896305 | 1:200 |
| CD45R | His24 | PE | BD Biosciences | 554881 | AB_395574 | 1:500 |
| CD45RA | OX33 | FITC | BD Biosciences | 554883 | AB_395576 | 1:200 |
| CD45RA | OX33 | APC-Cy7 | BD Biosciences | 561624 | AB_10893004 | 1:200 |
| TCR αβ | R7.3 | PerCP | BD Biosciences | 557019 | AB_396538 | 1:500 |
| IgM | MARM-4 | Pacific Blue | Home-made | / | / | 1:200 |
| IgD | MARD-4/3 | AF647 | Home-made | / | / | 1:200 |
| CD4 | OX35 | PE-Cy7 | BD Biosciences | 561578 | AB_10715836 | 1:500 |
| CD25 | OX39 | BV421 | BD Biosciences | 565608 | AB_2739303 | 1:200 |
| CD25 | OX39 | APC | Home-made | / | / | 1:500 |
| CD8 | OX8 | APC | Home-made | / | / | 1:500 |
| CD8 | OX8 | AF488 | Home-made | / | / | 1:500 |
| FoxP3 | FJK-16S | PE | Invitrogen / eBioscience | 12-5773-82 | AB_465936 | 1:50 |
| FoxP3 | FJK-16S | APC | Invitrogen / eBioscience | 17-5773-82 | AB_469457 | 1:50 |
| ICOS (CD278) | C398.4A | APC | BioLegend | 313510 | AB_416334 |  |
| Viability Dye | N/A | eFluor 450 | Invitrogen / eBioscience |  | N/A | 1:1000 |
| Viability Dye | N/A | eFluor 506 | Invitrogen / eBioscience |  | N/A | 1:1000 |
| 123count eBeads™ | N/A | Multi-fluorescent | Invitrogen / eBioscience | 01-1234-42 | AB_2574163 | 10 µl / test |
